# *In silico* design of a T-cell epitope vaccine candidate for parasitic helminth infection

**DOI:** 10.1101/859017

**Authors:** Ayat Zawawi, Ruth Forman, Hannah Smith, Iris Mair, Murtala Jibril, Munirah Albaqshi, Andrew Brass, Jeremy P. Derrick, Kathryn J. Else

## Abstract

*Trichuris trichiura* is a parasite that infects 500 million people worldwide, leading to colitis, growth retardation and Trichuris dysentery syndrome. There are no licensed vaccines available to prevent *Trichuris* infection and current treatments are of limited efficacy. *Trichuris* infections are linked to poverty, reducing children’s educational performance and the economic productivity of adults. We employed a systematic, multi-stage process to identify a candidate vaccine against trichuriasis based on the incorporation of selected T cell epitopes into virus-like particles. We conducted a systematic review to identify the most appropriate *in silico* prediction tools to predict histocompatibility complex class II (MHC-II) molecule T-cell epitopes. These tools were used to identify candidate MHC-II epitopes from predicted ORFs in the *Trichuris* genome, selected using inclusion and exclusion criteria. Selected epitopes were incorporated into Hepatitis B core antigen virus-like particles (VLPs). A combined VLP vaccine containing four *Trichuris* MHC-II T-cell epitopes stimulated dendritic cells and macrophages to produce pro-inflammatory and anti-inflammatory cytokines. The VLPs were internalized and co-localized in the antigen presenting cell lysosomes. Upon challenge infection, mice vaccinated with the VLPs+T-cell epitopes showed a significantly reduced worm burden, and mounted *Trichuris*-specific IgM and IgG2c antibody responses. The protection of mice by VLPs+T-cell epitopes was characterised by the production of mesenteric lymph node (MLN)-derived Th2 cytokines and goblet cell hyperplasia. Collectively our data establishes that a combination of *in silico* genome-based CD4+ T cell epitope prediction, combined with VLP delivery, offers a promising pipeline for the development of an effective, safe and affordable helminth vaccine.

**Author Summary:** The soil transmitted helminth *Trichuris trichiura* is a major parasite in developing countries; development of a comprehensive vaccine has been elusive. Here we used a systematic approach based on *in silico* identification of MHC-II T cell epitopes from genome sequences, their incorporation into a virus-like particle (VLP), characterization of the assemblies and testing in an *in vivo* murine infection model. Animals vaccinated with a preparation of four different VLP-antigen fusions showed significant reductions in intestinal worm burdens and associated antibody responses consistent with protection. The results suggest that a pipeline based on *in silico* prediction of potent MHC-II T cell epitopes, followed by incorporation into VLPs, could be a strategy which enables rapid translation into a vaccine against *Trichuris trichiura*.

## Introduction

Trichuriasis, caused by the whipworm *Trichuris trichiura*, is one of the most widespread soil-transmitted helminths (STH) in the world (1). Global mass drug administration (MDA) programmes are being implemented, but cure rates are low, repeated treatments are costly and may prevent the development of acquired immunity. Further, the existence of drug-resistant parasites is a constant concern (2–4). Using the mouse model of human trichuriasis, *Trichuris muris* excretory/secretory (ES) products (5), ES fractions (6), extracellular vesicles (EVs) (7), and, more recently, *T. muris* whey acidic protein (8) in the context of the adjuvant alum, have shown considerable potential in a number of pre-clinical protection trials. Despite these successes, developing a vaccine based on native antigens is associated with many manufacturing challenges, including cost, time consumption, difficulties in purifying large quantities of worm antigens and control over differences between batches (9, 10). The advent of the genome era has provided alternative strategies for vaccine development (11). For example, the reverse vaccinology (RV) approach combines genome information with immunological and bioinformatics tools to overcome some of the limitations of conventional methods of screening vaccine candidates (12–14).

These observations, combined with the economic challenges of producing a low-cost vaccine, prompted us to examine the potential of using virus-like particles (VLPs) as a scaffold for the presentation of predicted *Trichuris* MHC-II epitopes. The Hepatitis B core protein (HBc) has been widely used as a VLP: it forms stable self-assemblies, which can accommodate T cell epitopes, is cheap to produce and is safe for human use (15, 16). Structural analysis has shown that each HBc monomer forms helical structures, which protrude as spikes from the capsid (17). Antigens, in the form of short T cell epitopes or whole globular domains, can be inserted at the tip of each spike. Designing vaccines based on ‘multi-stage’ antigens which are expressed at different stages of infection have shown promising results against several complex pathogens, such as *M. tuberculosis* and *Plasmodium* (18, 19). Further, CD4+ Th2 cells play essential roles in the development of protective immunity against *Trichuris* spp (20).The principal objective of this study was therefore to develop a novel MHC II T-cell epitope-based vaccine predicted from multi-stage *Trichuris* proteins, which will induce Th2 protective immunity. To achieve this aim, first, a systematic review was performed to select the optimal MHC class II *in silico* prediction tools. Second, potential *Trichuris* MHC-II T-cell epitope vaccine candidates obtained from the *Trichuris* genome were identified using the selected *in silico* prediction tool. Third, these epitopes were produced in a commercially viable manner by fusing epitopes in the hepatitis B core antigen (HBc-Ag) virus-like particle (VLP) vaccine delivery system. These VLPs+T-cell epitope vaccine candidates were then tested *in vitro* for their ability to activate antigen presenting cells (APCs). Finally, *in vivo* experiments were conducted to test the protective capacity of VLPs expressing different *Trichuris* T-cell epitopes *in vivo* using *T. muris* infection of mice. Collectively the results of this research represent the first significant progress towards identifying a novel, epitope-based vaccine for trichuriasis.

## Results

### The IEDB and NetMHC-II 2.2 tools exhibited similarly high levels of sensitivity for predicting epitopes with strong affinities

Based on a list of search terms (Supplementary Table S1) used on Google and other websites (Supplementary Table S2), 88 servers that predict T-cell epitopes based on MHC class I and II binding were identified (Supplementary Table S3). Of these 88, only 48 tools which could predict MHC-II epitopes were identified. Since our primary focus was MHC-II T-cell epitope prediction, only tools with that functionality were further evaluated using the inclusion criteria (Fig 1A). Five tools met the inclusion criteria: IEDB, SYPEITHI, NetMHC-II 2.2, Rankpep and ProPred. These tools were then scanned for the ability to predict MHC-II T-cell epitopes for two mouse alleles, I-Ab and I-Ad. The ProPred tool was excluded from further analysis because it only predicted HLA-DR binding sites. The four epitope prediction tools that met all the selection criteria were subsequently evaluated using an epitope training set to calculate the sensitivity with which they could predict MHC II T-cell epitopes. Using the epitope training set (Supplementary Table S4), it was observed that IEDB and NetMHC-II 2.2 tools had high levels of sensitivity (~78.00%) for predicting MHC-II T-cell binding epitopes with high affinities, while Rankpep and SYFPETHI exhibited low sensitivity (10.61% and 8.33%, respectively). Collectively, the data indicate that the best prediction tools across all MHC-II T-cell prediction servers considered in this study are IEDB and NetMHC-II 2.2.

**Fig 1.**
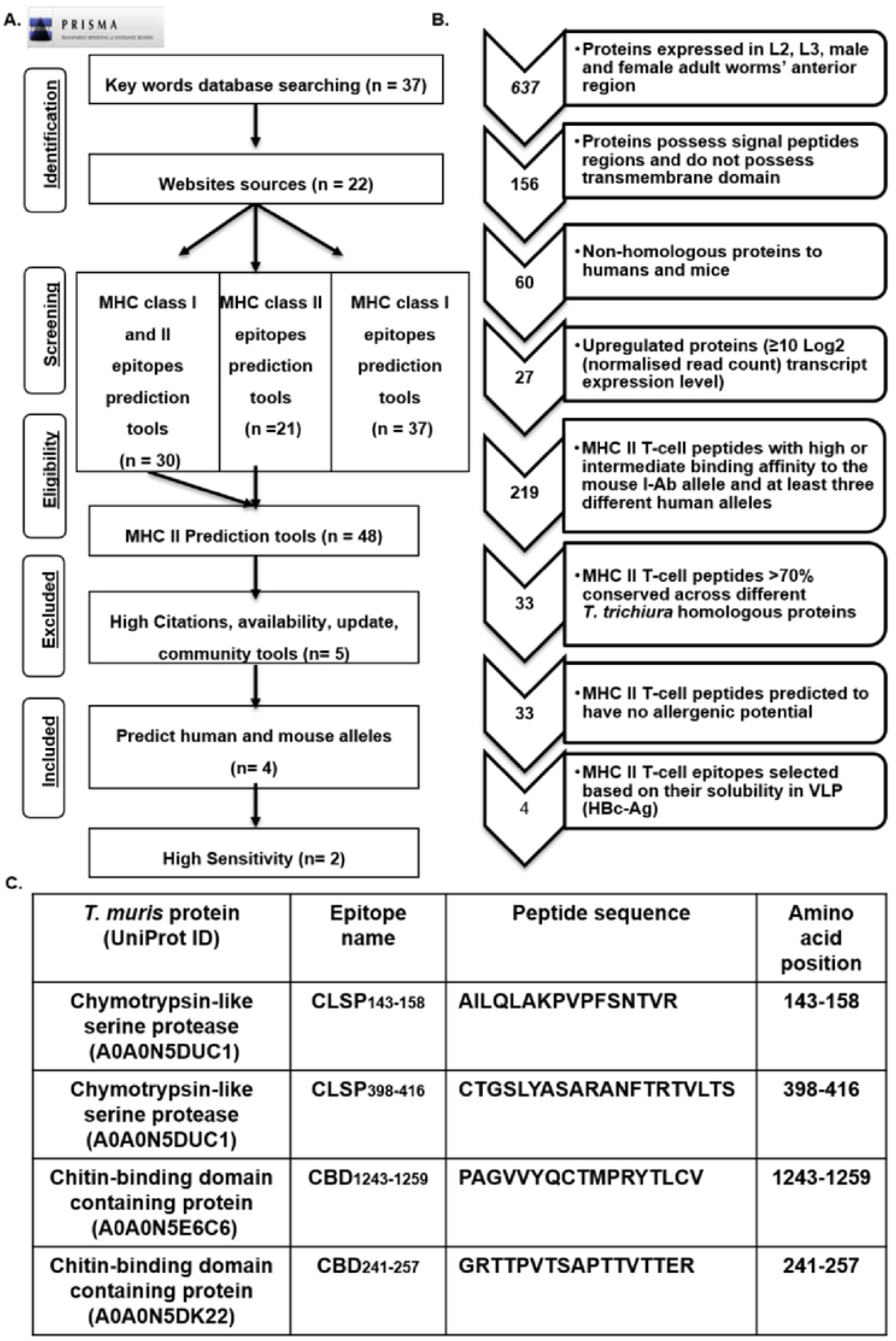
(A) PRISMA flow diagram of the systematic review screening process for restricted MHC-II T-cell bioinformatics prediction tools. A list of search terms was used in Google website and other website sources to screen for MHC class I and II *in silico* prediction tools. Aspects relating to the number of citations (>200), number of publications, online availability, last update and community were used as inclusion/exclusion criteria to select the top MHC-II *in silico* prediction tools. n= number of key words, websites sources and *in silico* prediction tools. (B) The flow diagram of reverse vaccinology approach to identify potential vaccine candidates (MHC-II T-cell epitopes) from the *T. muris* genome. The number on the arrow represents the number of proteins or epitopes selected for the next step. (C) List of 4 *Trichuris* MHC-II T-cell epitopes which have potential as vaccine candidates.

### Data acquisition and identification of potential vaccine component proteins

The stichosome, which forms the majority of the whipworm anterior region, is thought to release ES products which trigger the host immune response and induce host immunity after infection (7, 21). Research conducted by Dixon et al., (22) and Else et al., (23), showed that antibody recognition of high molecular weight proteins in both *T. muris* adult and larval ES correlated with resistance to *T. muris* infection. To increase the chances of the identification of antigens which induce a strong immune response, secreted and surface-exposed proteins associated with the anterior region of L2, L3, male and female adult worms were selected for this study (24).

Predicted ORFs from the 85-Mb genome (∼11,004 protein-encoding genes) of *T. muris* and the 73-Mb genome (∼9650 protein-coding genes) of *T. trichiura* (24) were scanned for potential vaccine candidates; 637 proteins were selected. From this subset, only proteins that possess signal peptides and did not exhibit transmembrane domains were included, reducing the total to a sample of 156 proteins. Full-length sequences of the *Trichuris* proteins were obtained from the Universal Protein Resource (UniProt) database in FASTA format http://www.uniprot.org/.

### Elimination of closely homologous mouse and human proteins

To eliminate potential autoimmune reactions when tested in mice and humans, the 156-protein subset was checked to determine homology with human and mouse proteins using the basic local alignment search tool (BLAST). All proteins with any degree of homology with humans or mice were excluded, leaving 60 candidate proteins. Of these, only upregulated proteins in the anterior region of L2, L3 and adult worms were selected based on their transcript expression level. This criterion was based on high-throughput transcriptome data generated from the RNA of *T. muris* and Gene Ontology (GO) term enrichment analyses, a transcriptional upregulation in a particular protein refers to a ≥10 Log2 (normalised read count) transcript expression level (24). Implementing this criterion, 27 proteins were selected for MHC-II T-cell epitope prediction (Fig 1B).

### Prediction of *Trichuris* MHC-II T-cell binding epitopes

All 27 selected proteins were screened to predict T-cell MHC class II epitopes using the IEDB (consensus method) prediction tool (25). The analysis was carried out to predict the binding affinity to the MHC class II allele I-Ab mouse strain and the 11 most prevalent human class II HLA allele supertypes (25–28). The consensus method was used to select only those peptides with a low median percentile rank according to three different prediction methods to reduce the chance of failure during prediction. To cover common global alleles, peptides were selected based on their ability to bind to at least three different human alleles.

### Conservation and allergens

To assess how well the predicted MHC-II T-cell peptides were conserved within the *T. trichiura* genome, the IEDB conservancy analysis tool was used. This tool calculates the degree of conservancy (i.e. similarity) of a peptide within a specified protein sequence (29). Only peptides that were >70% conserved with at least one homologous *T. trichiura* protein were selected for further analysis. Of the 219 MHC-II T-cell peptides, only 33 met these criteria. The 33 MHC-II peptides were then assessed for the prediction of IgE epitopes and allergenic potential using the AllerTOP v.2.0 server http://www.ddg-pharmfac.net/AllerTOP/ (30). The final set of 10 *Trichuris* MHC-II T-cell epitopes containing 33 overlapping peptides were predicted to have no allergenic potential. The 10 epitopes were further triaged to a final four epitopes based on solubility once expressed on HBc-Ag VLPs. A flow diagram summarising the approach used is shown in Fig 1B, with the final 4 *Trichuris* MHC II T-cell epitopes detailed in Fig 1C.

### Purification and assembly of VLPs expressing *Trichuris* T-cell epitopes

Four HBc-Ag fusion proteins were designed, incorporating each predicted T-cell epitope into the major immunodominant region. The construct also included a Strep tag at the C-terminus, for affinity purification (Supplementary Figure S1). A second round of purification using size exclusion chromatography was performed to produce a homologous population of assembled VLPs. HBc-Ag preparations expressing *Trichuris* MHC-II T-cell epitopes were pure by SDS PAGE (Supplementry Figure S1 B-E). The endotoxin levels in all purified VLPs used in this study were <0.2 endotoxin units (EU) as assessed using the ELISA-based endotoxin detection assay (Data not shown).

### VLPs expressing *Trichuris* T-cell epitopes induced the production of proinflammatory cytokines *in vitro* and were internalised and co-localized in APCs

BMDCs stimulated with different VLPs, irrespective of whether they include T-cell epitopes, produced high levels of IL-6 (Fig 2A) and TNF-α (B) at levels equivalent to LPS and ES-stimulated BMDCs. Similiarly, all VLPs activated BMDMs, inducing the secretion of high levels of both TNF-α (Fig. 2E), and IL-6 (F). To visualise VLP internalisation by APCs, BMDCs and BMDMs were stimulated with or without fluorescein-conjugated VLPs. Images of individual cells taken by merging the brightfield (BF) and FITC (green) channels demonstrated that the fluorescein-conjugated VLPs were internalised by both APCs (Fig 2C & G). To confirm that the VLPs were co-localized within the APC lysosome and not the cell surface, BMDCs and BMDMs were stained with the lysosome-specific LysoTracker dye. Cells were then subsequently stimulated with or without fluorescein-conjugated VLPs. Merged images of the brightfield (BF), FITC (green), and Lysotracker (red) channels revealed that the fluorescein-conjugated VLPs were accumulated in the lysosome compartment of the BMDCs (Fig 2D) and BMDMs (Fig 2H).

**Fig 2.**
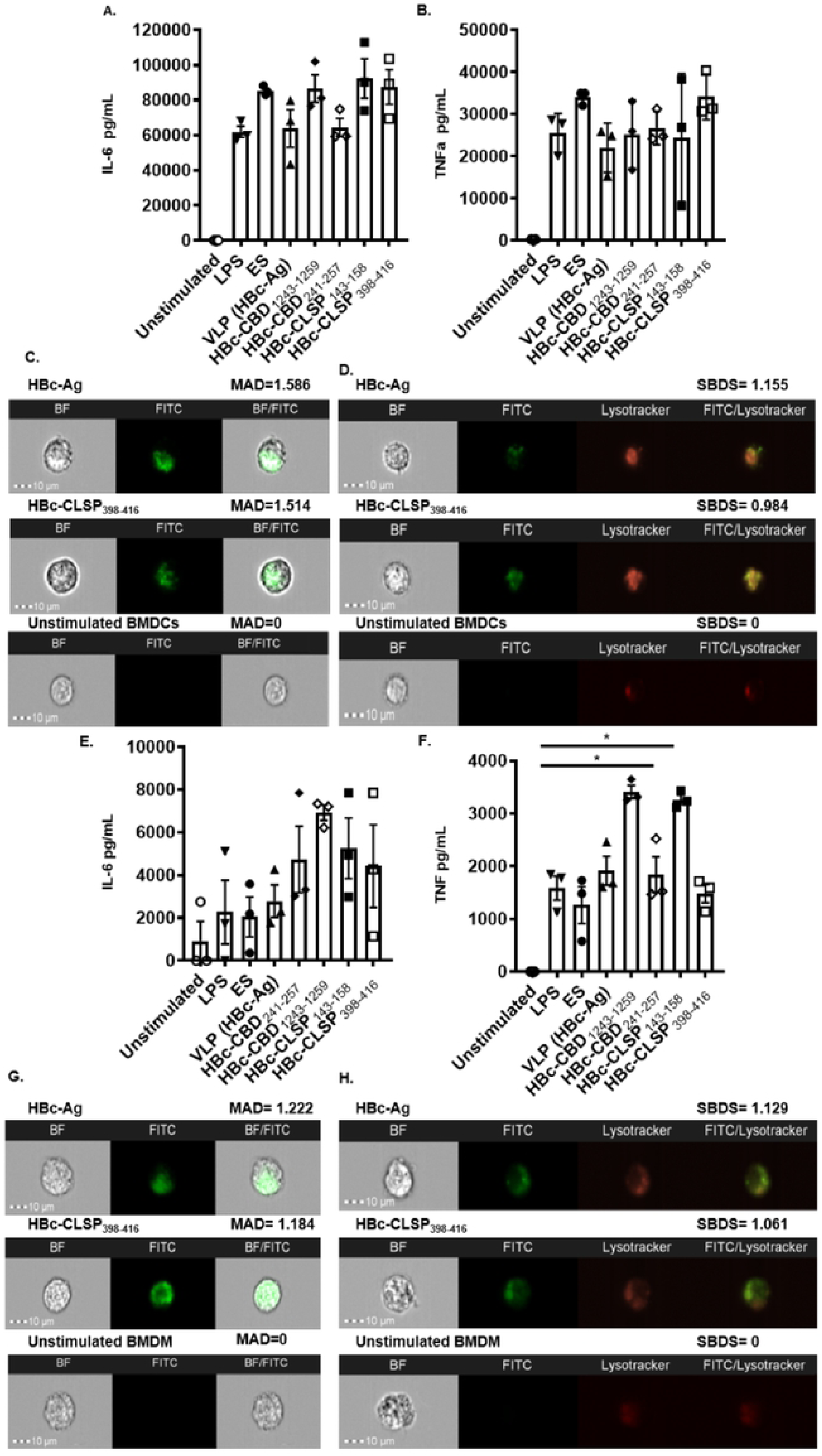
(A) Inflammatory cytokines production by mouse bone marrow-derived DCs (BMDCs) and mouse bone marrow-derived macrophages (BMDMs) (B) in response to VLPs. BMDCs and BMDMs at 1×10^6^/ml were stimulated *in vitro* with 10 µg/ml VLPs (HBc-Ag, HBc-H_112-128_, HBc-CBD_1243-1259,_ HBc-CBD_241-257_, HBc-CLSP_143-158_ and HBc-CLSP_398-416_) and with 50 µg/ml ES and 0.1 µg/ml LPS as positive controls. Unstimulated BMDCs and BMDMs served as negative controls. Supernatants were harvested after 24 hours for IL-6 and TNF α cytokine analyses measured by CBA. The bars represent mean ± SEM. Statistical analyses were carried out using the Kruskal-Wallis test (multiple comparisons). Significant differences between groups are represented by (*P≤0.05) with a line. Chart bars represent BMDCs and BMDMs grown from three individual mice from one representative experiment of two separate experiments. (C) Representative images of fluorescein-conjugated VLPs internalisation in the BMDCs and BMDMs (G). BMDCs and BMDMs at 1×10^6^/ml were incubated with 10 μg/ml fluorescein-conjugated VLP (HBc-Ag and HBc-CLSP_398-416_) for 24 hours. As a negative control, unstimulated BMDCs and BMDMs were examined. Cell internalisation was determined by Amnis ImageStreamX cytometer compared to unstimulated BMDCs and BMDMs. Images shown, from left to right, show individual Brightfield images (BF) in the white channel, fluorescent-labelled stimulus (FITC) in the green channel and the combination of both BF/FITC merged channels. The internalisation mean absolute deviation (MAD) is included above its images. The positive MAD value represents internalisation, and negative values represent poor internalisation. (D) Representative images of fluorescein-conjugated VLPs co-localization in the BMDCs and BMDMs (H). BMDCs and BMDMs at 1×10^6^/ml were stained with Lysotracker to visualise the cellular lysosome compartment and subsequently stimulated with 10 μg/ml fluorescein-conjugated VLP (HBc-Ag, and HBc-CLSP_398-416_) for 24 hours. As a negative control, unstimulated BMDCs and BMDMs were examined. Intracellular co-localization was determined by Amnis ImageStreamX cytometer. Images shown, from left to right, show individual Brightfield images (BF) in the white channel, fluorescent-labelled stimulus (FITC) in the green channel, stained lysosome (Lysotracker) in the red channel and the combination of both FITC/ Lysotracker merged channels. The Similarity bright detail score (SBDS) from the IDEAS quantitative co-localization analysis is included above its image. SBDS values around 1 represent co-localization, and 0 values represent poor co-localization. Scale bars represent 10 μm.

### Immunization of mice with VLPs expressing *Trichuris* T-cell epitopes induced a significant reduction in worm burden following challenge infection

The protective capacity of the 4 T cell epitopes was assessed in the *T. muris* – mouse accelerated expulsion model (31) (Fig 3A). Mice vaccinated with pre-mixed four VLPs+T-cell epitopes showed a statistically significant reduction in worm burden by day 14 p.i. compared to the native VLP (HBc-Ag; (P<0.01) and PBS (P<0.05) immunised groups (Fig 3B). In comparison, ES/Alum immunised mice harboured no parasites (Fig 3C).

**Fig 3.**
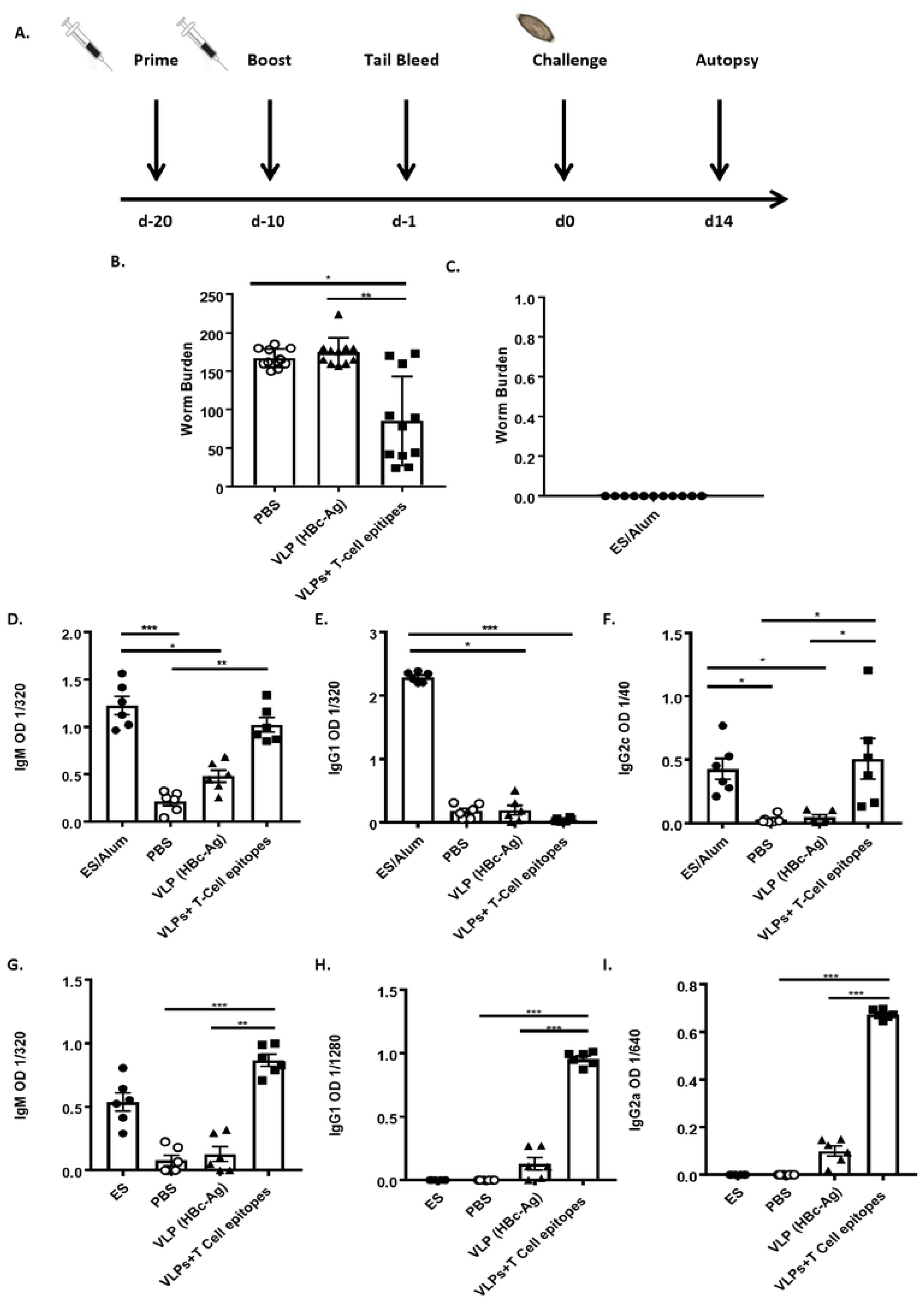
Experimental schedule, worm burden, parasite-specific and VLP-specific antibodies levels of mice vaccinated and challenged with a *T. muris* infection. 6-week old male C57BL/6 mice were vaccinated subcutaneously with 50 μg of pre-mixed of VLPs+T-cell epitopes (HBc-CBD_1243-1259_, HBc-CBD_241-257_, HBc-CLSP_143-158_, and HBc-CLSP_398-416_), 50 μg of VLP (HBc-Ag), or PBS on day −20 and boosted on day −10. Mice were infected by oral gavage with approximately 150 infective *T. muris* eggs on day 0 and sacrificed at d14 post-infection. (B) Comparison of worm burden at day 14 post-infection. (C) Worm burden in mice vaccinated with 50 μg of ES. n= 11 mice per group. The results presented are from two separated experiments pooled together. Day 14 p.i. sera were titrated against *T. muris* ES antigens to assess parasite-specific IgM (D) IgG1 (E), and IgG2c (F) levels in VLPs+T-cell epitopes, VLP (HBc-Ag), PBS and in ES vaccinated mice by ELISA (reading at 405 nm). Day 14 p.i. sera were titrated against premixed VLPs+ T-cell epitopes antigens to assess VLPs+ T-cell epitopes-specific IgM (G) IgG1 (H), and IgG2c (I) levels in VLPs+T-cell epitopes, VLP (HBc-Ag), PBS and in ES vaccinated mice. Statistical analyses were carried out using Kruskal-Wallis test (multiple comparisons). Significant differences between groups are represented by (*P≤0.05, **P≤0.01, ***P≤0.001, ****P≤0.0001) with a line. Results are shown as mean ± SEM. n= 6 mice per group. This experiment was repeated two times, and the ELISA results shown here are representative of the two experiments.

### Immunization of VLPs expressing *Trichuris* T-cell epitopes induced humoral immunity following challenge infection

To evaluate *T. muris*-specific serum antibody responses induced by vaccination with VLPs+T-cell epitopes, parasite-specific IgM, IgG1, and IgG2c serum antibody levels were determined at d14 p.i. Following vaccination and infection, VLPs+T-cell epitopes and ES/Alum vaccinated mice had statistically significant higher levels of parasite-specific IgM (Fig 3D) and IgG2c (F) compared to the control PBS and native VLP (HBc-Ag) injected mice. However, high levels of parasite-specific IgG1 were only detected in the serum of mice vaccinated with ES/Alum following *Trichuris* infection (Fig 3E). There was no or very low levels of parasite-specific IgM, IgG1, and IgG2c detected in the serum of native VLP (HBc-Ag) and PBS/alum injected mice at day 14 p.i., as shown in Fig. (3D-F). Similarly, statistically significant higher levels of VLPs+T-cell epitopes-specific IgM (Fig 3G), IgG1 (H) and IgG2c (I) were produced following *Trichuris* infection of mice vaccinated with VLPs+T-cell epitopes, compared to mice given native VLP (HBc-Ag) or PBS. Notably, mice vaccinated with ES/Alum also produced high levels of VLPs+T-cell epitopes-specific IgM following *Trichuris* infection, as shown in Fig 3 (G).

### Immunisation of mice with VLPs expressing *Trichuris* T-cell epitopes induces a mixed Th1/Th2 immune response following challenge infection

To analyse the cellular immune responses at the primary site of adaptive immune cell activation following *T. muris* infection (32), MLN cells of mice vaccinated with VLPs+T-cell epitopes were re-stimulated *in vitro* with *Trichuris* ES. Supernatants were assayed for Th2 cytokines (IL-4, IL-5, IL-9 and IL-13), Th1/Th17 cytokines (IL-2, IFN-γ and IL-17), proinflammatory cytokines (IL-6 and TNF-α), and the anti-inflammatory cytokine (IL-10) production by CBA (Fig 4 and Supplementary Figure S2). Levels reached significance versus control vaccinated mice for IL-5 (Fig 4A; ES vaccinated mice) and IFN-γ (Fig 4D; VLP+T cell epitope vaccinated mice) with non-significant increases in IL-9 and IL-13 (Fig 4B, 4C). MLN cells from all vaccinated mouse groups also produced IL-4, IL-6, IL-10, IL-17 and TNF upon their re-stimulation *in vitro* (Supplementary Figure S2 A-E). Collectively these data support the *in vivo* immunogenicity of the novel VLP+ T cell epitope vaccine, evidencing the presence of a mixed Th1/Th2 immune response and are in keeping with the worm expulsion and antibody data. Proximal colon goblet cells were quantified in mice vaccinated with ES/Alum, PBS, VLP (HBc-Ag) and VLPs+T-cell epitopes day 14 post *T. muris* infection. (Fig 4E). Interestingly, VLPs+T-cell epitopes and ES/Alum vaccinated mice exhibited a significantly elevated goblet cell hyperplasia (P<0.05) compared to mice injected with PBS or native VLP (HBc-Ag) (Fig 4F).

**Fig 4.**
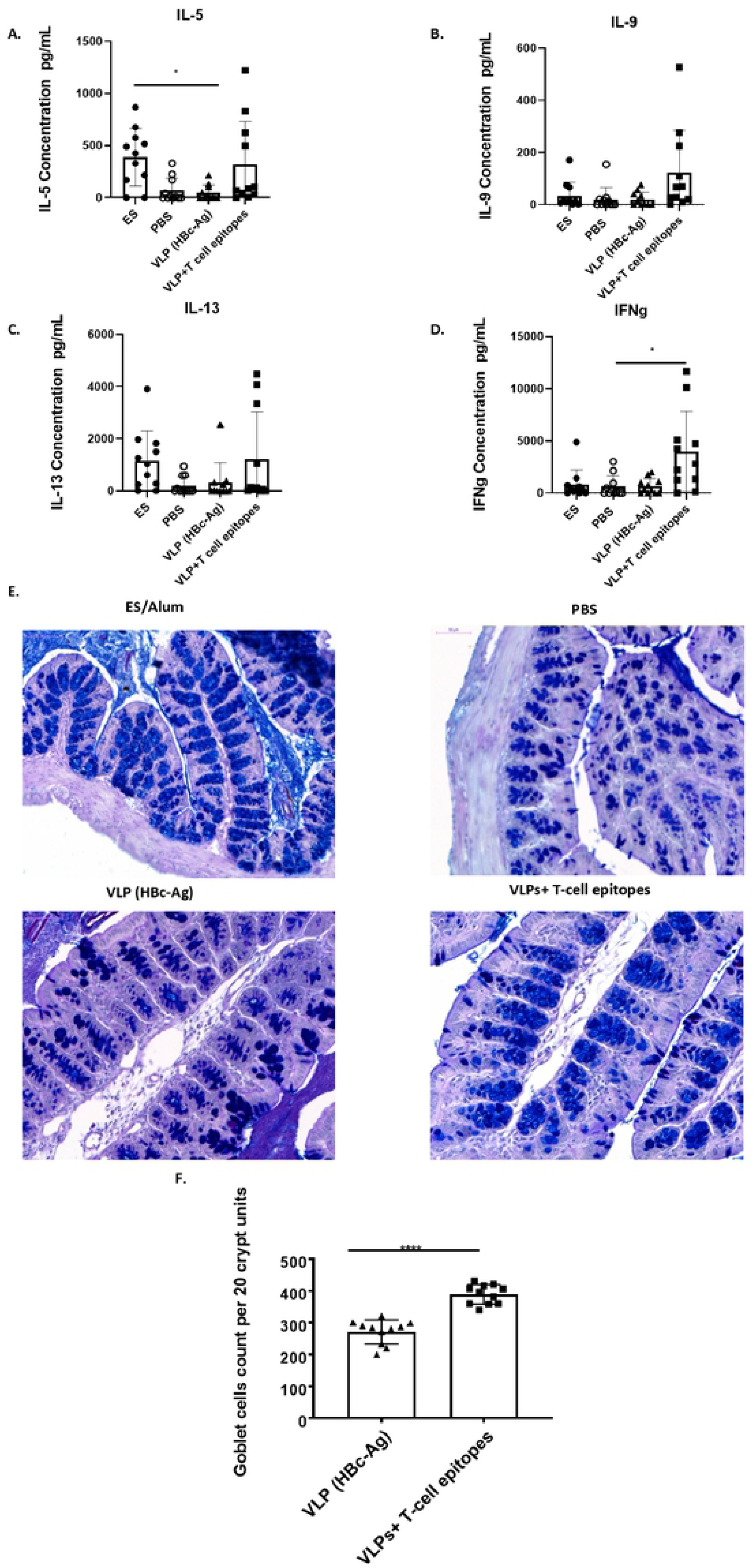
Cytokine productions by mesenteric lymph node cells and quantification of goblet cell numbers in mice vaccinated with VLPs+T-cell epitopes and control mice. (A) MLNs from mice vaccinated with 50 μg of pre-mixed of VLPs+T-cell epitopes epitopes (HBc-CBD_1243-1259_, HBc-CBD_241-257_, HBc-CLSP_143-158_ and HBc-CLSP_398-416_), 50 μg of VLP (HBc-Ag), 50 µg ES/Alum, or PBS on day −20 and boosted on day −10 were stimulated at 5×10^6^/ml with 50 µg/ml *T. muris* ES. After 48 hours of stimulation, cell culture supernatants were harvested and assayed by cytometric bead array for IL-5 (A), IL-9 (B), IL-13 (C), IFN-γ (D) production. (E) Representative photographs of gut sections stained with Periodic acid–Schiff staining (PAS). The slides were scanned, and the goblet cells were quantified using Panoramic Viewer software. All photographs were taken with 20X magnification. (F) Goblet cells were quantified by counting the number of alcian blue-stained cells per crypt unit in three fields of view from each section and are shown as mean cell numbers per 20 crypt units (cu) ±SEM. Statistical analyses were carried out using the Kruskal-Wallis test (multiple comparisons). Significant differences between groups are represented by (*P≤0.05, ****P≤0.0001) with a line. Results are shown as mean ± SEM. n= 11 mice per group. The results presented are from two separated experiment pooled together.

## Discussion

*Trichuris trichiura* is one of the most common human STH parasites and remains a major health concern for humans worldwide (1). A number of pre-clinical vaccines against trichuriasis have been reported, containing whole *Trichuris* antigens or fractions (6, 7, 22). However, developing a vaccine for *Trichuris* based on native antigens has several limitations (10). An alternative strategy embraces the use of informatics to predict and assess MHC-II T cell epitopes using criteria to maximize their *in vivo* protective potential. The fast-growing application of VLP-based vaccines against several parasites, including *Plasmodium* spp (33, 34), *Leishmania infantum* (35), *Toxoplasma gondii* (36), *Trichinella spiralis* (37), and *Clonorchis sinensis* (38) prompted our interest in developing a VLP-based vaccine against trichuriasis.

### Identification of novel *Trichuris* MHC-II T-cell epitopes as promising vaccine candidates

Initiation of an antigen-specific immune response requires presentation of antigenic peptides to CD4+ T-cells in the context of MHC Class II molecules (39). Selection of appropriate MHC Class II binding peptides is therefore a critical first step in developing an epitope-based vaccine. There are more than 80 computer-based prediction tools for identifying peptides that bind to MHC class I and II molecules, but not all are equivalent; some epitope prediction tools may fail to predict all significant epitopes (40). The performance of *in silico* prediction tools is affected by several factors. For example, MHC-II CD4+ T-cell epitope prediction tools have much lower accuracy than MHC-I tools because the MHC-I binding groove is closed, while the MHC-II groove is open at both ends (40, 41). Also, the use of a limited training dataset to evaluate the tools may affect performance (25, 42, 43). Further, the low performance of MHC class II prediction tools may not only be due to poor algorithm performance; the genetic diversity of human populations presents additional challenges (44). Thus, choosing the best bioinformatics tool to predict MHC class II T-cell epitopes is critical when designing epitope-based vaccines (45, 46). This study presents a systematic review of existing MHC-II restricted T-cell epitope prediction tools and evaluated four tools in order to establish the most appropriate bioinformatics tools currently available for predicting *Trichuris* MHC-II T-cell epitopes.

The IEDB and NetMHC-II 2.2 tools achieved similarly high levels of sensitivity of predicting binding epitopes with high affinities, while Rankpep and SYFPETHI exhibited low sensitivity. Each tool applies different methods of prediction; the IEDB tool uses a quantitative consensus method that combines the strengths of various methods (25); the NetMHC-II 2.2, another quantitative tool, uses an NN-align algorithm and weight matrix (28); and both SYFPEITHI (47) and Rankpep (48) are qualitative tools that use motif PSSMs. All the tools were ‘user-friendly’ but SYFPETHI was limited in its coverage of mouse and human MHC-II alleles. The IEDB tool has features that the other three do not, including a browser to input protein sequence formats in a National Centre for Biotechnology Information (NCBI) database, seven different prediction methods and an easy method of downloading the prediction output into an Excel spreadsheet. Several studies have compared the performance of different MHC class II peptide binding prediction tools (49–51), but the comparison presented in this study is different for two main reasons. First, the MHC-II prediction tools were selected in a systematic way using inclusion/exclusion criteria. Second, a unique dataset consisting of a new ‘test’ dataset which has not been used to build or evaluate IEDB was used. For example, Zhao and Sher (2018) evaluated the MHC-II prediction tools hosted on the IEDB analysis resource server using newly available, untested data of both synthetic and naturally processed epitopes. Among the 18 predictors that were benchmarked, NetMHC-II outperformed all other tools, including NetMHCIIpan and the consensus method for both MHC class I and class II predictions (52). Furthermore, Andreatta et al. (2018) created an automated platform to benchmark six commonly used MHC class II peptide binding prediction tools using 59 datasets that were newly entered into the IEDB database and had not yet been made public to prevent biased assessment of the available prediction tools. Their evaluation suggested that NetMHCIIpan is currently the most accurate tool, followed by NN-align and the IEDB (consensus) tool (53). Despite differences in the datasets used for comparison, these studies agree with the comparison study conducted in our study, which found that the IEDB (consensus) and NETMHC-II 2.2 (ANN) tools are among the best MHC-II prediction tools. However, NetMHCIIpan could not be included in this study because it did not meet the inclusion criteria. Collectively, we recommend the use of IEDB and NETMHC-II 2.2 prediction tools in any MHC class II epitope prediction study to reduce the experimental cost of identifying epitopes. However, the output of these tools needs to be carefully evaluated *in vitro* and *in vivo* before they are used to bring an epitope-based vaccine to trial.

Since the advent of immunoinformatics tools for prediction of antigenic epitopes and protein analysis, several VLP-based vaccines have been engineered to carry foreign antigens and have proven to be highly immunogenic (54). To our knowledge, the data presented in this study is the first to identify novel *Trichuris* MHC-II T-cell epitopes as potential vaccine antigen candidates. The framework used here to identify potential epitope vaccine candidates within the *T. muris* and *T. trichiura* genome could be used in the future to identify potential vaccine candidates for other parasite species. The final set of *Trichuris* MHC-II T-cell epitope vaccine candidates were derived from chitin-binding domain-containing proteins and chymotrypsin-like serine proteases (Fig 1C).

Chitin-binding domains genes are highly expressed in different life stages in many nematodes including the parasites *T. trichiura*, *Ascaris lumbricoides* and *Ancylostoma ceylanicum* and the free-living nematode *Caenorhabditis elegans*, (55–58). In particular, these proteins are thought to be associated with eggshell formation and early development at the single-cell stage (59, 60). Furthermore, given that, in addition to *T. muris*, more than 40 other helminth species express high levels of chymotrypsin-like serine proteases, these proteins may also be promising vaccine candidates for other helminths (24, 61). Chymotrypsin-like serine proteases are thought to play central roles in either the invasion process or modulation of the host immune response to enhance parasite survival (62–65). In addition, numerous publications, using pre-clinical models of parasitic infection, have noted that protective immunity can be consistently achieved using helminth protease molecules (66). For example, vaccinating BALB/c mice with the whole recombinant serine protease of *T. spiralis* prior to challenge infection led to a reduction in worm burden and induced a mixed Th1/Th2 immune responses (67–69). Furthermore, Shears et al. (6) showed that vaccinating mice with the *T. muris* ES fraction containing serine proteases induced high parasite-specific antibody responses. Remarkably all the proteins selected in this study have been identified within the most immunogenic fractions of *T. muris* ES following vaccination of mice (6).

### Identification of a novel VLP-based vaccine against trichuriasis

All VLP recombinant proteins stimulated a non-specific inflammatory response characterized by the secretion of high levels of proinflammatory cytokines. Furthermore, by identification of intracellular co-localization with lysosomes, VLPs were shown to be taken up by both BMDC and BMDMs. These results are consistent with those of Serradell et al. (70) and Wahl-Jensen et al. (71) who examined the activation of APCs in response to VLPs. For example, Serradell et al. developed a (VLP-HA/VSP-G) vaccine composed of an influenza virus expressing influenza virus hemagglutinin (HA) antigen, co-expressed with the extracellular region of *Giardia lamblia* variant-specific surface proteins (VSPs) as an adjuvant. The recombinant protein (VLP-HA/VSP-G) induced the secretion of high levels of TNF-α, IL-10, and IL-6 cytokines *in vitro*. They also showed that oral vaccination with the recombinant protein protected mice from influenza infection and generated protective humoral and cellular immunity. These results suggest that the VLPs are well placed to act as delivery systems to drive immune responses. Further, the data raises the exciting prospect of modifying the VLPs using adhesion molecules and/or cytokines co-displayed on the VLP surface, in order to target the epitopes to specific APC subsets, thus enhancing the activation of antigen-specific T-cells (72).

Remarkably, upon challenge with *T. muris* infection, mice vaccinated with 50 μg of pre-mixed VLP+T-cell epitopes (HBc-CBD_1243-1259_, HBc-CBD_241-257_, HBc-CLSP_143-158_, and HBc-CLSP_398-416_), in the absence of any additional adjuvant, showed a significantly reduced worm burden. Parasite-specific IgM and IgG2c were detected in the sera of mice vaccinated with the VLPs+T-cell epitopes. Levels were equivalent to the control ES in alum-vaccinated mice, and significantly higher than control vaccinated and infected mice. These results suggest that these VLPs+T-cell epitopes are antigenic and can boost antibody response sufficiently to recognize specific small peptides in the *T. muris* ES. Importantly, analysis of the serum from immunised mice showed that vaccination with the VLPs+T-cell epitopes elicited high levels of IgM, IgG1 and IgG2c to the VLP+T cell epitope recombinant protein pool (HBc-CBD_1243-1259_, HBc-CBD_241-257_, HBc-CLSP_143-158_, and HBc-CLSP_398-416_) with limited recognition of the native VLP (HBc-Ag) protein. In keeping with these data, the malaria VLP-based vaccine (Malarivax) (73–75) composed of HBc-Ag expressing *P. falciparum* T cell and B-cell epitopes identified from the circumsporozoite protein developed long-lasting immunity, eliciting a CD4+ T cell immune response and is currently undergoing a clinical trial (15, 76, 77). The results of these studies support some critical insights into VLP-based vaccines, including confirming that an HBc virus-like particle, in particular, is an excellent delivery system for developing potential vaccine candidates for parasites.

A proportion of mice vaccinated with VLP+T-cell epitopes produced detectable levels of MLN-derived Th2 cytokines IL-5, IL-9, and IL-13 in response to re-stimulation *in vitro* with *Trichuris* ES. The Th1 cytokine IFN-γ was also significantly elevated above levels detected in control mice. These data indicate that vaccine-induced protective immunity is characterized by a mixed Th1/Th2 immune response, as has been previously reported (21, 78). Similarly, Gu et al (78), reported that the protective immunity to *Trichinella spiralis* infection induced by vaccination with CD4+ T cell epitopes was associated with both Th1 and Th2 cytokines. Whilst the mechanism by which vaccination protects mice from *T. muris* infection remains unclear, this study reveals that vaccination of mice with VLPs+T-cell epitopes or ES/Alum promoted a marked goblet cell hyperplasia (79). Goblet cells, and the mucins they produce, have been implicated in Th2-mediated defence in mice resistant to a primary *T. muris* infection (80, 81). However, it remains to be determined whether the goblet cell hyperplasia seen here in vaccinated mice is simply a Th2 correlate or is functionally important in the protection observed. In summary, the current study describes the development and efficacy of a novel epitope-based vaccine against trichuriasis. VLPs expressing different *Trichuris* MHC-II T-cell epitopes, predicted from chitin-binding domain containing proteins and chymotrypsin-like serine proteases, were shown to promote protective immunity *in vivo*. Collectively, given the right combination of immunoinformatics and immunogenicity screening tools, epitope-based vaccines will undoubtedly limit the cost and effort associated with bringing a *Trichuris* vaccine to trial.

## Methods

### Search strategy

A protocol was designed to identify the bioinformatics tools that can predict MHC-II T-cell epitopes in accordance with the well-defined Preferred Reporting Items for Systematic Reviews and Meta-Analyses (PRISMA) (82). The search was limited to the English language and used the search terms shown in (Supplementary Table S1). A list of the search terms was first used in December 2015 on Google and other websites (Supplementary Table S2) to screen for MHC class I and II *in silico* prediction tools. The number of citations (>200), number of publications (>200), online availability, last update and community were considered to determine whether the tools would be selected for further analysis. The number of publications and citations for each tool were obtained from Google Scholar. In the case of duplicate citations, the highest number of citations was used. A flow diagram of the systematic review screening process for MHC-II T-cell bioinformatics prediction tools is shown in Fig. 1A.

### Construction of an epitope training set

To evaluate the performance of the tools, a literature search for MHC-II CD4+ T-cell peptide binding datasets was performed using the search terms ‘universal T-cell epitopes’ and ‘MHC class II T-cell epitopes’ on Google Scholar in December 2015. These datasets included epitopes with publicly available sequences that the literature had experimentally validated to be immunogenic in mice. Because training sets of peptides were used for the development of epitope prediction tools, care was taken to exclude the training datasets used for tool development (83–85).

In order to find the optimal set of epitopes, two different sets of epitopes were analysed. The first set was composed of publicly available series of peptide sequences from 15 different proteins that can bind to MHC class II molecules. The second set included one untested protein of *T. muris*, which served as a control to evaluate the performance of the tools. Collectively, the training dataset was composed of 145 epitopes from 16 different proteins (Supplementary Table S4).

### Prediction of *Trichuris* MHC-II T-cell epitopes

Full-length sequences of proteins containing T-cell epitopes were obtained from the Universal Protein Resource (UniProt) database http://www.uniprot.org in FASTA format. The Immune Epitope Database (IEDB), NetMHC-II 2.2, Rankpep and SYFPEITHI tools were used to predict T-cell epitopes for mouse strains with I-Ab and I-Ad mouse alleles. Each tool applied a different prediction method and scoring system to generate the predictions output. For instance, the IEDB tool uses the consensus method for prediction, which combines the NN-align, SMM-align, combinatorial library and Sturniolo methods (25), while the NetMHC 2.2 tool uses ANNs (28). The median percentile rank (%) of the three prediction methods was used to generate the rank for the consensus method. A small numbered percentile rank (%) indicates that a peptide has a high binding affinity to MHC class II alleles (25).The output of the IEDB and NetMHC 2.2 tools (the binding affinities) were expressed as half-maximal inhibitory concentration IC50 (nM) values. Epitopes that bind with an affinity of <50 nM are considered to have high affinity, those that bind with <500 nM have an intermediate affinity and those that bind with <5000 nM have a low affinity (28). All epitopes with high or intermediate affinity are considered ‘true binders’, while epitopes with low affinity are considered ‘nonbinders’. No known T-cell epitope has an IC50 value of >5000 nM (25). The scoring system of the SYFPEITHI prediction tool depends on whether peptide amino acids are frequently occur in anchor positions. Optimal anchor residues are given the value 15 and scores of −1 or −3 points are given to amino acids that have a negative effect on epitopes’ binding ability at a certain sequence position. Epitopes that bind strongly are among the top 2% of all peptides predicted in 80% of all predictions results (47). The Rankpep tool uses position-specific scoring matrices (PSSMs) to predict MHC-II T-cell epitopes (48). A high peptide score percentage indicates that the epitope is likely to bind to the set of aligned peptides that bind to a given MHC-II molecule (48). The peptide lengths in all the resulting sets were based on the complex of 15 mer core region peptides of MHC class II molecules.

### Evaluation and statistical analysis

Using the training set of epitopes, the performance of the four MHC-II epitope prediction tools, selected through our inclusion/exclusion criteria, were assessed as weak, intermediate or high binders. The prediction results were classified into two categories, true positive (TP) and false negative (FN), based on the threshold values (86). In addition, the evaluation assessed sensitivity (TP/ [TP+FN]). Nonparametric Spearman correlation and Bland-Altman analyses were performed to show the relationship and agreement between the scores derived from the NetMHC-II 2.2 and IEDB tools. The level of significance was set at p < 0.05 for the correlation test.

### Mice and Parasites

6–8 weeks male C57BL/6 mice (Envigo) were fed autoclaved food and water and were maintained under specific pathogen-free conditions. Parasite maintenance, ES collection from adult *T. muris* worms and the method used for infection and evaluation of worm burden were carried out as described previously (87).

### Ethics Statement

All animal experiments were approved by the University of Manchester Animal Welfare and Ethical Review Board and performed under the regulation of the Home Office Scientific Procedures Act (1986), and the Home Office approved licence 70/8127.

### Plasmid construction

The coding sequence for native VLP (HBc-Ag) was engineered with BamHI + EcoRI sites to allow insertion of peptide antigen sequences into the major immunodominant region (MIR). The entire HBc-Ag coding sequence was inserted into the pET17b expression vector between the NdeI (CATATG) and XhoI (CTCGAG) restriction sites. The insertion of each MHC-II T-cell epitope into the MIR was obtained by annealing the relevant oligonucleotide primers (Supplementary Table S5) with BamHI + EcoRI restriction sites and ligation into BamHI/EcoRI cut HBc-Ag in the pET-17b vector. Constructs containing MHC-II T-cell epitopes were confirmed by DNA sequencing.

### Production and purification of VLP recombinant proteins

The recombinant plasmids were transformed into ClearColi-BL21 (DE3) electrocompetent cells (Lucigen) by heat-shock transformation. The bacterial culture was grown in LB media containing 100 μg/mL ampicillin and incubated at 37°C for 14 hours in a shaker incubator. The transformed cells at a starting optical density (0.1 OD600) were inoculated into LB liquid media supplemented with 100 μg/mL ampicillin for 3-4 hours until the optical density at 595 nm (OD600) reached 0.6. Isopropyl-b-D-thiogalactopyranoside (IPTG) was added to the culture to a final concentration of 0.4 mM, and the culture was grown by continuous shaking for 12-16 hours at 16°C. The cells were then harvested by centrifugation, resuspended and sonicated in Strep-Tag washing buffer (100 mM Tris-HCl, 150 mM NaCl, 1 mM EDTA) contaning 1 tablet of cOmplete TM EDTA-free protease inhibitor cocktail (Sigma-Aldrich) per 50 mls of the resuspended cells along with 5 µg/ml DNase I (Sigma-Aldrich). The cell suspension was then disrupted by ultra-sonication on ice using Banddelinuw 3200 (Sonoplus) amplitude (Am) in 35% for 5 sec on and 10 sec off for 5-8 minutes. The supernatant containing soluble recombinant protein was harvested following centrifugation at 18,900 x g for 40 mins at 4°C using Sorvall RC Plus with the Fiber-Lite F21 8×50y rotor. Finally, the soluble supernatant was filtered through 0.22 µm pore size filters. After filtration, Strep(II)-tag proteins were purified using affinity column chromatography using a StrepTrap™ column prepacked with StrepTactin Sepharose by following the manufacturer’s protocol (GE Healthcare). The VLP recombinant proteins were further purified by size exclusion chromotogrophy (Superose 6, 10/300 GL;GE Healthcare). The level of endotoxin in all the purified VLPs was measured with an ELISA-based endotoxin detection assay (Hyglos) following the manufacturer’s protocol.

### SDS-PAGE

The VLP recombinant protein samples were subjected to 10% SDS-PAGE and subsequently characterized by TEM as described previously [20]. Briefly, the VLP recombinant proteins were separated on 10% polyacrylamide gels and stained with instant blue protein stain (Expedeon).

### *In vitro* stimulation of BMDCs and BMDMs

BMDCs were harvested from mice into RPMI 1640© (Sigma-Aldrich) supplemented with 5% heat-inactivated FCS (Hyclone, Logon, UT), 100 μg /mL penicillin and 100 μg/mL streptomycin (Sigma-Aldrich), 2 mM L-Glutamine (Sigma-Aldrich), 50 µM β-mercaptoethanol (Gibco) and 20 ng/ml granulocyte-macrophage colony-stimulating factor (GM-CSF) (eBioscience). BMDMs were harvested from mice into modified Eagle′s Medium (DMEM) – high glucose (Sigma-Aldrich) supplemented with 5% heat-inactivated FCS (Hyclone, Logon, UT), 100 μg/mL penicillin and 100 μg/mL streptomycin (Sigma-Aldrich), Eagle′s minimum essential medium (MEM) non-essential amino acid solution (Sigma-Aldrich), and 0.1 mg/ml macrophage colony-stimulating factor (M-CSF) (eBioscience). BMDMs and BMDCs on day 8 were collected at 1×10^6^/ml and incubated with 10 µg/ml VLP recombinant protein for overnight at 37°C. ES at a final concentration 50 µg/ml and lipopolysaccharide (LPS) at final concentration 0.1 µg/ml were used as positive controls, and untreated cells were used as a negative control. After 24 hours, the supernatants were harvested separately and stored at −20°C until analysed by LEGENDplex for cytokine content according to the manufacturer’s instructions.

### Fluorescein-conjugated VLP internalization and localization in APCs

BMDCs and BMMs on day 8 were collected at 1×10^6^/ml and incubated with 10 µg/ml with fluorescein-conjugated VLP separately (HBc-CBD_1243-1259,_ HBc-CBD_241-257,_ HBc-CLSP_143-158_ and HBc-CLSP_398-416_) for overnight at 37°C in 5% CO_2_. Unstimulated BMDCs and BMMs were used as negative controls. Next day, cells were incubated with 50 mM LysoTracker dye obtained from (Invitrogen) for 45 min prior to harvesting to visualise the lysosomal localisation of fluorescein-conjugated VLPs by ImageStreamX cytometry. A Brightfield-1 filter was employed to image dendritic cells and macrophages, a fluorescein isothiocyanate (FITC-488 nm) filter to image fluorescein-conjugated VLPs, and an (APC-592 nm) filter to image lysosomes. The data were analysed using IDEAS software version 6.2.187.0. The degree of co-localization was measured by the Bright Detail Similarity (BDS-R3) on a cell-by-cell basis. A Bright Detail Similarity value of 1.0 indicates a high degree of similarity between two images in the same spatial location (correlated) and a value around 0 has no significant similarity (uncorrelated).

### Immunization schedule and challenge infection

Mice were divided randomly into 4 groups: mice were inoculated s.c with overnight ES antigens emulsified with an equal volume of Alum-an aluminum salt adjuvant (88) obtained from Thermo Scientific to achieve a vaccine dose of 100 μg ES in 100 μl Alum as a positive control. The negative group was inoculated s.c with 200 µl of sterile PBS. Alternatively, mice were vaccinated with 200 µl of 50 μg VLP (HBc-Ag). The vaccinated mouse group was administrated with 200 µl of 50 μg pre-mixed VLPs+T-cell epitopes (HBc-CBD_1243-1259,_ HBc-CBD_241-257,_ HBc-CLSP_143-158_ and HBc-CLSP_398-416_). All experimental groups were vaccinated in the scruff of the neck at (day −20) and were boosted with the same dose at (day −10) in the same site. At day 10 post vaccination, mice were challenged orally with a high dose of *T. muris* (200 eggs) per mouse. Mice were sacrificed 14 days after challenge infection by an overdose of CO_2_.

### Ag-specific antibody detection in the serum

Blood samples were collected immediately from mice by cardiac puncture and left at room temperature to clot. Parasite-specific and VLP recombinant protein-specific antibodies (IgM, IgG1 and IgG2c) were determined in sera by enzyme-linked immunosorbent assay (ELISA) as previously described (89). Briefly, 96-well plates were coated with 50 μg/well of the overnight *T. muris* E/S antigen at 5 µg/ml or with 50 μg/well of the purified VLP recombinant protein in 0.5 M carbonate-bicarbonate buffer for overnight at 4°C. The plates were washed and then blocked with 3% BSA (Sigma-Aldrich) in PBS Tween-20 (PBS-T20) (0.05% Tween 20, Sigma-Aldrich) for 1 h at 37 °C. After washing, diluted sera were added and incubated for 1 h at 37 °C. Antibody responses were detected using the Biotinylated rat anti-mouse IgM, IgG1 and IgG2a/c (BD Biosciences). After washing, streptavidin peroxidase (Sigma-Aldrich) was added to the plates and incubated for 1 hour at room temperature. The TMB ELISA substrate (3, 3’, 5, 5’-tetramethylbenzidine-Thermo) was used to develop color and stopped with 0.003% (H_2_SO_4_). The optical density at 450 nm with a reference of 570 nm was measured with a Dynex MRX11 plate reader (DYNEX Technologies) at 450 nm with a reference of 570 nm.

### *In vitro* stimulation of MLN

MLNs were harvested from mice at autopsy and dissociated in RPMI-1640 medium (Sigma-Aldrich) supplemented with 10% foetal calf serum (FCS), 1% L-Glutamine, and 1% penicillin/streptomycin (all Invitrogen Ltd., Paisley, UK). 5×10^6^/ml MLN cells were re-stimulated *in vitro* in 96-well plates with 50 μg/ml *T. muris* E/S antigens and with 10 µg/ml pre-mixed VLPs+T-cell epitopes proteins (HBc-CBD_1243-1259_, HBc-CBD_241-257_, HBc-CLSP_143-158_, and HBc-CLSP_398-416_) at 37 °C for 48 hours. Cytokine concentration was quantified in the supernatants by cytometric bead array (CBA) according to the manufacturer’s instructions from the Mouse/Rat Soluble Protein Flex Set System kit (BD Biosciences Pharmingen, Oxford, UK). CBA data was analysed on a MACSQuant analyser with MACSQuantify™ Software and the CBA analysis software package (BD Biosciences).

### Histology

At autopsy, colonic tissue samples were removed and fixed at room temperature for overnight in 10% neutral buffered formalin, prior to storage in 70% EtOh and processing and embedding in paraffin wax. 5 μm thick serial sections were cut on a Microrn HM325 microtome (Microm International, Germany), de-waxed in citroclear and rehydrated prior to periodic acid–Schiff staining. Stained slides of proximal colon sections were scanned using 3D Histech Pannoramic 250 flash slide scanner. Photographs of the sections were taken at 100X magnification using panoramic viewer version 1.15.4 software. The number of goblet cells was determined as the total number of PAS-positive cells in 60 randomly selected crypts in three fields of view from each section.

### Statistics

Statistical analyses were performed using Graph Pad Prism, version 7.00 software. In all tests, P≤0.05 were considered statistically significant and were determined using the Kruskal–Wallis non-parametric ANOVA for comparing multiple groups.

## Acknowledgments

We thank the University of Manchester Histology, Bio-imaging and Flow Cytometry Core Facility for facilitating the image analyses.

## Supporting Information Legends

**S1 Fig. The HBc-antigen fusion recombinant protein purification and assembly. Left: Chromatogram of the VLP recombinant proteins HBc-Ag (A), HBc-CBD_241-257_ (C), HBc-CLSP_143-158_, (D), and HBc-CLSP_398-416_ (E) purified using SEC.** Separation was carried out on a Superose 6, 10/300 GL (GE Healthcare), at a flow rate of 0.2 ml/min with 100 mM Tris-HCl, 150 mM NaCl, 1 mM EDTAatpH 8. VLP recombinant proteins were eluted in 0.5 ml fraction and a sample of SEC the elution peak was visualised by SDS-PAGE. Vertical red lines indicate the elution volume of mass standards, from left to right: 2MDa, 670kDa. Right: Purification visualised by SDS-PAGE (10% gel) analysis stained with Coomassie blue. M: protein marker (Precision Plus protein standard, unstained, Bio-Rad); Lane 1: VLP recombinant protein after Strep-Tag affinity purification using the StrepTrap HP, and Lane 2: VLP recombinant protein after SEC purification. The arrow represents VLP recombinant protein monomer with an expected molecular mass of ~22 kDa.

**S2 Fig. Cytokine productions by mesenteric lymph node cells and quantification of goblet cell numbers in mice vaccinated with VLPs+T-cell epitopes and control mice.** (A) MLNs from mice vaccinated with 50 μg of pre-mixed of VLPs+T-cell epitopes epitopes (HBc-CBD_1243-1259_, HBc-CBD_241-257_, HBc-CLSP_143-158_ and HBc-CLSP_398-416_), 50 μg of VLP (HBc-Ag), 50 µg ES/Alum, or PBS on day −20 and boosted on day −10 were stimulated at 5×10^6^/ml with 50 µg/ml *T. muris* ES. After 48 hours of stimulation, cell culture supernatants were harvested and assayed by cytometric bead array for IL-4 (A), IL-6 (B), IL-10 (C), IL-17 (D), and TNF (E) production. Statistical analyses were carried out using the Kruskal-Wallis test (multiple comparisons). Significant differences between groups are represented by (*P≤0.05) with a line. Results are shown as mean ± SEM. n= 11 mice per group. The results presented are from two separated experiment pooled together.

**S1 Table. List of Keywords used to screen for MHC I and II *in silico* prediction tools.**

**S2 Table. List of MHC class I and II and HLA binders peptides *in silico* prediction tools.**

**S3 Table. MHC-II T-cell epitopes dataset used to evaluate the MHC-II *in silico* prediction tools.**

**S4 Table. List of MHC-II T-cell epitope sequences and primers used to amplify the target**.

